# Widespread natural variation of DNA methylation within angiosperms

**DOI:** 10.1101/045880

**Authors:** Chad E. Niederhuth, Adam J. Bewick, Lexiang Ji, Magdy S. Alabady, Kyung Do Kim, Qing Li, Nicholas A. Rohr, Aditi Rambani, John M. Burke, Josh A. Udall, Chiedozie Egesi, Jeremy Schmutz, Jane Grimwood, Scott A. Jackson, Nathan M. Springer, Robert J. Schmitz

## Abstract

To understand the variation in genomic patterning of DNA methylation we compared methylomes of 34 diverse angiosperm species. By analyzing whole-genome bisulfite sequencing data in a phylogenetic context it becomes clear that there is extensive variation throughout angiosperms in gene body DNA methylation, euchromatic silencing of transposons and repeats, as well as silencing of heterochromatic transposons. The Brassicaceae have reduced CHG methylation levels and also reduced or loss of CG gene body methylation. The Poaceae are characterized by a lack or reduction of heterochromatic CHH methylation and enrichment of CHH methylation in genic regions. Reduced CHH methylation levels are found in clonally propagated species, suggesting that these methods of propagation may alter the epigenomic landscape over time. These results show that DNA methylation patterns are broadly a reflection of the evolutionary and life histories of plant species.

## Introduction

Biological diversity is established at multiple levels. Historically this has focused on studying the contribution of genetic variation. However, epigenetic variations manifested in the form of DNA methylation (Vaughn, Tanurdzic et al. 2007, Eichten, Swanson-Wagner et al. 2011, Schmitz, Schultz et al. 2013), histones and histone modifications (Filleton, Chuffart et al. 2015), which together make up the epigenome, may also contribute to biological diversity. These components are integral to proper regulation of many aspects of the genome; including chromatin structure, transposon silencing, regulation of gene expression, and recombination(Colome-Tatche, Cortijo et al. 2012, Jones 2012, Schmitz and Zhang 2014). Significant amounts of epigenomic diversity are explained by genetic variation (Bell, Pai et al. 2011, Eichten, Swanson-Wagner et al. 2011, Eichten, Briskine et al. 2013, Regulski, Lu et al. 2013, Schmitz, He et al. 2013, Schmitz, Schultz et al. 2013, Dubin, Zhang et al. 2015), however, a large portion remains unexplained and in some cases these variants arise independently of genetic variation and are thus defined as “epigenetic” (Becker, Hagmann et al. 2011, Eichten, Swanson-Wagner et al. 2011, Schmitz, Schultz et al. 2011, Eichten, Briskine et al. 2013, Regulski, Lu et al. 2013, Schmitz, He et al. 2013). Moreover, epigenetic variants are heritable and also lead to phenotypic variation (Cubas, Vincent et al. 1999, Thompson, Tor et al. 1999, Manning, Tor et al. 2006, Silveira, Trontin et al. 2013). To date, most studies of epigenomic variation in plants are based on a handful of model systems. Current knowledge is in particular based upon studies in *Arabidopsis thaliana*, which is tolerant to significant reductions in DNA methylation, a feature that enabled the discovery of many of the underlying mechanisms. However, *A. thaliana*, has a particularly compact genome that is not fully reflective of angiosperm diversity (Arabidopsis Genome 2000, Flowers and Purugganan 2008). The extent of natural variation of mechanisms that lead to epigenomic variation in plants, such as cytosine DNA methylation, is unknown and understanding this diversity is important to understanding the potential of epigenetic variation to contribute to phenotypic variation (Lane, Niederhuth et al. 2014).

In plants, cytosine methylation occurs in three sequence contexts; CG, CHG, and CHH (H=A, T, or C), are under control by distinct mechanisms (Niederhuth and Schmitz 2014). Methylation at CG (mCG) and CHG (mCHG) sites is typically symmetrical across the Watson and Crick strands (Finnegan, Genger et al. 1998). mCG is maintained by the methyltransferase MET1, which is recruited to hemi-methylated CG sites and methylates the opposing strand (Finnegan, Peacock et al. 1996, Bostick, Kim et al. 2007), whereas mCHG is maintained by the plant specific CHROMOMETHYLASE 3 (CMT3) (Lindroth, Cao et al. 2001), and is strongly associated with dimethylation of lysine 9 on histone 3 (H3K9me2) (Du, Zhong et al. 2012). The BAH and CHROMO domains of CMT3 bind to H3K9me2, leading to methylation of CHG sites (Du, Zhong et al. 2012). In turn, the histone methyltransferases KRYPTONITE (KYP), and Su(var)3-9 homologue 5 (SUVH5) and SUVH6 recognize methylated DNA and methylate H3K9 (Du, Johnson et al. 2014), leading to a self-reinforcing loop (Du, Johnson et al. 2015). Asymmetrical methylation of CHH sites (mCHH) is established and maintained by another member of the CMT family, CMT2 (Zemach, Kim et al. 2013, Stroud, Do et al. 2014). CMT2, like CMT3, also contains BAH and CHROMO domains and methylates CHH in H3K9me2 regions (Zemach, Kim et al. 2013, Stroud, Do et al. 2014).

Additionally, all three sequence contexts are methylated *de novo* via RNA-directed DNA methylation (RdDM) (Law and Jacobsen 2010). Short-interfering 24 nucleotide (nt) RNAs (siRNAs) guide the *de novo* methyltransferase DOMAINS REARRANGED METHYLTRANSFERASE 2 (DRM2) to target sites (Cao and Jacobsen 2002, Cao and Jacobsen 2002). The targets of CMT2 and RdDM are often complementary, as CMT2 in *A. thaliana* primarily methylate regions of deep heterochromatin, such as transposons bodies (Zemach, Kim et al. 2013). RdDM regions, on the other hand, often have the highest levels of mCHH methylation and primarily target the edges of transposons and the more recently identified mCHH islands (Gent, Ellis et al. 2013, Zemach, Kim et al. 2013, Stroud, Do et al. 2014). The mCHH islands in *Zea mays* are associated with upstream and downstream of more highly expressed genes where they may function to prevent transcription of neighboring transposons (Gent, Ellis et al. 2013, Li, Gent et al. 2015). The establishment, maintenance, and consequences of DNA methylation are therefore highly dependent upon the species and upon the particular context in which it is found.

Sequencing and array-based methods allow for studying methylation across entire genomes and within species (Vaughn, Tanurdzic et al. 2007, Schmitz, Schultz et al. 2011, Schmitz, Schultz et al. 2013, Stroud, Greenberg et al. 2013, Dubin, Zhang et al. 2015). Whole genome bisulfite sequencing (WGBS) is particularly powerful, as it reveals genome-wide single nucleotide resolution of DNA methylation (Cokus, Feng et al. 2008, Lister, O’Malley et al. 2008, Lister and Ecker 2009). WGBS has been used to sequence an increasing number of plant methylomes, ranging from model plants like *A. thaliana* (Cokus, Feng et al. 2008, Lister, O’Malley et al. 2008) to economically important crops like *Z. mays* (Eichten, Swanson-Wagner et al. 2011, Gent, Ellis et al. 2013, Regulski, Lu et al. 2013, Li, Eichten et al. 2014). This has enabled a new field of comparative epigenomics, which places methylation within an evolutionary context (Feng, Cokus et al. 2010, Zemach, McDaniel et al. 2010, Takuno and Gaut 2012, Takuno and Gaut 2013). The use of WGBS together with *de novo* transcript assemblies has provided an opportunity to monitor the changes in methylation of gene bodies among species (Takuno, Ran et al. 2016) but does not provide a full view of changes in the patterns of context-specific methylation at different types of genomic regions (Seymour, Koenig et al. 2014).

Here, we report a comparative epigenomics study of 34 angiosperms (flowering plants). Differences in mCG and mCHG are in part driven by repetitive DNA and genome size, whereas in the Brassicaceae there are reduced mCHG levels and reduction/losses of CG gene body methylation (gbM). The Poaceae are distinct from other lineages, having low mCHH levels and a lineage-specific distribution of mCHH in the genome. Additionally, species that have been clonally propagated often have low levels of mCHH. Although some features, such as mCHH islands, are found in all species, their association with effects on gene expression is not universal. The extensive variation found suggests that both genomic, life history, and mechanistic differences between species contribute to this variation.

## Results

### Genome-wide DNA methylation variation across angiosperms

We compared single-base resolution methylomes of 34 angiosperm species that have genome assemblies (Sato, Nakamura et al. 2008, van Bakel, Stout et al. 2011, Goodstein, Shu et al. 2012, Dohm, Minoche et al. 2014) (**Table S1**). MethylC-seq (Lister, O’Malley et al. 2008, Urich, Nery et al. 2015) was used to sequence 26 species and an additional eight species with previously published methylomes were downloaded and reanalyzed (Schmitz, Schultz et al. 2011, Amborella Genome 2013, Gent, Ellis et al. 2013, Schmitz, He et al. 2013, Stroud, Ding et al. 2013, Zhong, Fei et al. 2013, Seymour, Koenig et al. 2014). Different metrics were used to make comparisons at a whole-genome level. The genome-wide weighted methylation level (Schultz, Schmitz et al. 2012) combines data from the number of instances of methylated cytosine sites relative to all sequenced cytosine sites, giving a single value for each context that can be compared across species (Figure 1A–C). The proportion that each methylation context makes up of all methylation indicates the predominance of specific methylation pathways (Figure 1D). The per-site methylation level is the distribution of methylation levels at individual methylated sites and indicates within a population of cells, the proportion that are methylated (Figure 1E-1G). Symmetry is a comparison of per-site methylation levels at cytosines on the Watson versus the Crick strand for the symmetrical CG and CHG contexts (**Figure S1,2**). The asymmetry of CHG sites are further quantified (**Figure S3**)based on symmetric methylation levels previously determined from *A. thaliana cmt3* mutants (Bewick, Ji et al. 2016). Per-site methylation and symmetry provide information into how well methylation is maintained and how ubiquitously the sites are methylated across cell types within sequenced tissues (Willing, Rawat et al. 2015).

**Fig. 1.**
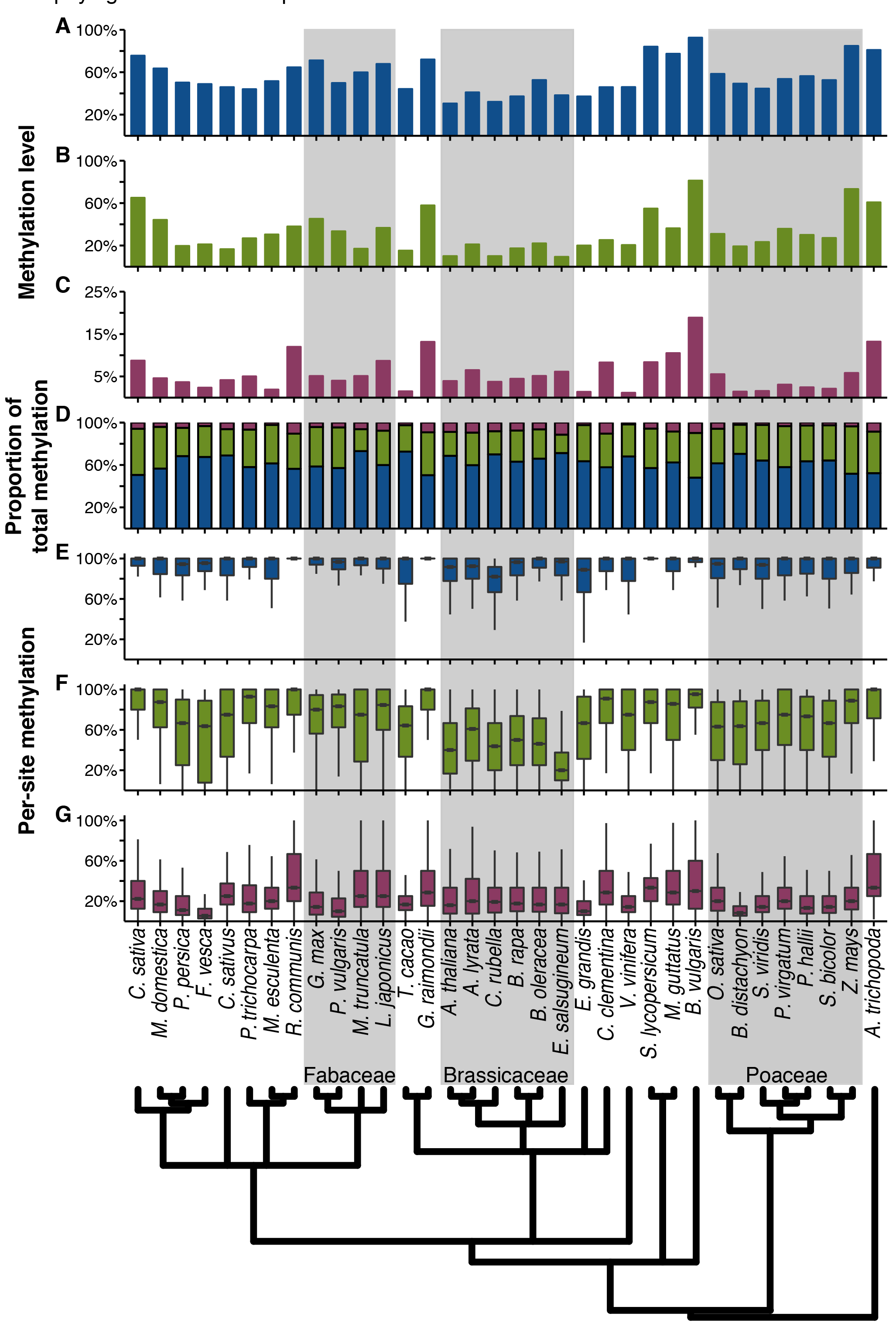
Genome-wide methylation levels for **(A)** mCG, **(B)** mCHG, **(C)** and mCHH. **(D)** Using the genome-wide methylation levels, the proportion that each context contributes towards the total methylation (mC) was calculated. **(E)** The distribution of per-site methylation levels for mCG, **(F)** mCHG, **(G)** and mCHH. Species are organized according to their phylogenetic relationship.

These data revealed that DNA methylation is a major base modification in flowering plant genomes. However, there is extensive variation between species, suggesting differences in activity amongst known DNA methylation pathways and potentially undiscovered DNA methylation pathways. Within each species, mCG had the highest levels of methylation genome-wide (Figure 1A, **Table S2**). Between species, levels ranged as much as three fold, from a low of ~30.5% in *A. thaliana* to a high of ~92.5% in *Beta vulgaris.*. Levels of mCHG varied as much as ~8 fold between species, from only ~9.3% in *Eutrema salsugineum* to ~81.2% in *B. vulgaris*, (Figure 1B, **Table S2**). mCHH levels were universally the lowest, but also the most variable with as much as an ~16 fold difference. The highest being ~18.8% is in *B. vulgaris.* This was unusually high, as 85% of species had less than 10% mCHH and half had less than 5% mCHH (Figure 1C, **Table S2**). The lowest mCHH level was found in *Vitis vinifera* with only ~1.1% mCHH. mCG is the most predominant type of methylation making up the largest proportion of the total methylation in all examined species (Figure 1D). *B. vulgaris* was a notable outlier, having the highest levels of methylation in all contexts, and having particularly high mCHH levels. Multiple factors may be contributing to the differences between species observed, ranging from genome size and architecture, to differences in the activity of DNA methylation targeting pathways.

We examined these methylomes in a phylogenetic framework, which led to several novel findings regarding the evolution of DNA methylation pathways across flowering plants. The Brassicaceae, which includes *A. thaliana*, all have the lowest levels of per-site mCHG methylation (Figure 1F). Furthermore, symmetrical mCHG sites have a wider range of methylation levels and increased asymmetry, whereas, non-Brassicaceae species have very highly methylated symmetrical sites (**Figure S2,3B**), suggesting that the CMT3 pathway is less effective in Brassicaceae genomes. This is further evidenced by *E. salsugineum*, with the lowest mCHG levels (Figure 1b), is a natural *cmt3* mutant whereas *CMT3* is under relaxed selection in other Brassicaceae (Bewick, Ji et al. 2016, Bewick, Niederhuth et al. 2016). Methylation of CG sites is also less well maintained in the Brassicaceae, with *Capsella rubella* showing the lowest levels of per-site mCG methylation (Figure 1E, **S1**).

Within the Fabaceae, *Glycine max* and *Phaseolus vulgaris*, which are in the same lineage, show considerably lower per-site mCHH levels as compared to *Medicago truncatula* and *Lotus japonicus*, even though they have equivalent levels of genome-wide mCHH (Figure 1C,1G). The Poaceae, in general, have much lower levels of mCHH (~1.4−5.8%), both in terms of total methylation level and as a proportion of total methylated sites across the genome. Per-site mCHH level distributions varied, with species like *Brachypodium distachyon* having amongst the lowest of all species, whereas others like *O. sativa* and *Z. mays* have levels comparable to *A. thaliana.* In *Z. mays*, CMT2 has been lost (Zemach, Kim et al. 2013), and it may be that in other Poaceae, mCHH pathways are less efficient even though CMT2 is present. Collectively, these results indicate that different DNA methylation pathways may predominate in different lineages, with ensuing genome-wide consequences.

Several dicot species showed very low levels of mCHH (< 2%): *V. vinifera, Theobroma cacao, Manihot esculenta* (cassava), *Eucalyptus grandis.* No causal factor based on examined genomic features or examined methylation pathways was identified, however, these plants are commonly propagated via clonal methods (McKey, Elias et al. 2010). Amongst non-Poaceae species, the six lowest mCHH levels were found in species with histories of clonal propagation (**Figure S4**). Effects of micropropagation on methylation in *M. esculenta* using methylation-sensitive amplified polymorphisms have been observed before (Kitimu, Taylor et al. 2015), so has altered expression of methyltransferases due to micropropagation in *Fragaria* x *ananassa* (strawberry) (Chang, Zhang et al. 2009). If clonal propagation was responsible for low mCHH, we hypothesized that going through sexual reproduction might result in increased mCHH levels, as work in *A. thaliana* suggests that mCHH is reestablished during reproduction (Slotkin, Vaughn et al. 2009, Calarco, Borges et al. 2012). To test this, we examined a DNA methylome of a parental *M. esculenta* plant that previously undergone clonal propagation and a DNA methylome of its offspring that was germinated from seed. Additionally, the original *F. vesca* plant used for this study had been micropropagated for four generations. We germinated seeds from these plants, as they would have undergone sexual reproduction and examined these as well. Differences were slight, showing little substantial evidence of genome-wide changes in a single generation of sexual reproduction (**Figure S5**). As both of these results are based on one generation of sexual reproduction it may be that this is insufficient to fully observe any changes. This will require further studies of samples collected over multiple generations from matching lines that have been either clonally propagated or propagated through seed for numerous generations.

### Genome architecture of DNA methylation

DNA methylation is often associated with heterochromatin. Two factors can drive increases in genome size, whole genome duplication (WGD) events and increases in the copy number for repetitive elements. The majority of changes in genome size among the species we examined is due to changes in repeat content as the total gene number in these species only varies two-fold, whereas the genome size exhibits ~8.5 fold change. As genomes increase in size due to increased repeat content it is expected that DNA methylation levels will increase as well. This was tested using phylogenetic generalized least squares (PGLS) (Martins and Hansen 1997) which takes into account the phylogenetic relationship of species and modifies the slope and intercept of the regression line (**Table S3**). This methodology minimizes the influence of outlier species and the effects of having numerous closely related species of on the slope of the regression line. Phylogenetic relationships were inferred from a species tree constructed using 50 single copy loci for use in PGLS (**Figure S6**) (Duarte, Wall et al. 2010). Correlations from PGLS were corrected for the total number of PGLS tests conducted. Indeed, positive correlations were found between mCG (p-value = 2.9 × 10^−3^) and the strongest correlation for mCHG and genome size (p-value = 2.2 × 10^−6^) (Figure 2A). No correlation was observed between mCHH and genome size after multiple testing correction (p-value > 0.05) (Figure 2A). A relationship between genic methylation level and genome size has been previously reported (Takuno, Ran et al. 2016). We found that within coding sequences (CDS) methylation levels were correlated with genome size for both mCHG (p-value = 5 × 10^−6^) and mCHH (p-value = 1.4 × 10^−5^), but not for mCG (p-value > 0.18) (Figure 2B).

**Fig. 2.**
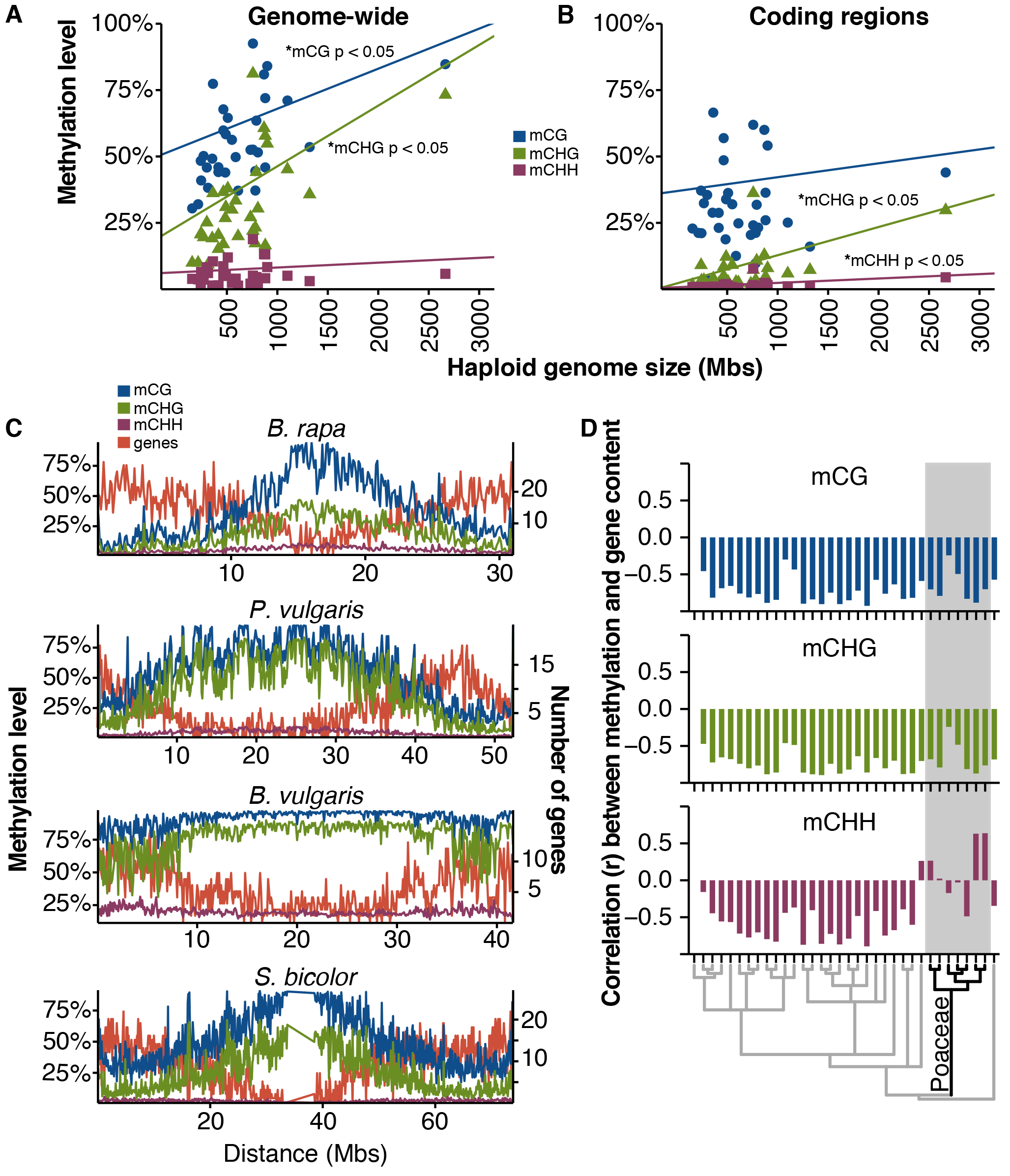
**(A)** Genome-wide methylation levels are correlated to genome size for mCG (blue) and mCHG (green), but not for mCHH (maroon) Significant relationships indicated. **(B)** Coding region (CDS) methylation levels is not correlated to genome size for mCG (blue), but is for mCHG (green), and mCHH (maroon). Significant relationships indicated. **(C)** Chromosome plots show the distribution of mCG (blue), mCHG (green), and mCHH (maroon) across the chromosome (100kb windows) in relationship to genes. **(D)** For each species, the correlation (Pearson’s correlation) in 100kb windows between gene number and mCG (blue), mCHG (green), and mCHH (maroon).

The highest levels of DNA methylation are typically found in centromeres and pericentromeric regions (Cokus, Feng et al. 2008, Lister, O’Malley et al. 2008, Seymour, Koenig et al. 2014). The distributions of methylation at chromosomal levels were examined in 100kb sliding windows (Figure 2C, **S7**). The number of genes per window was used as a proxy to differentiate euchromatin and heterochromatin. Both mCG and mCHG have negative correlations between methylation level and gene number, indicating that these two methylation types are mostly found in gene-poor heterochromatic regions (Figure 2D). Most species also show a negative correlation between mCHH and gene number, even in species with very low mCHH levels like *V. vinifera.* However, several Poaceae species show no correlation or even positive correlations between gene number and mCHH levels. Only two grass species showed negative correlations, *Setaria viridis* and *Panicum hallii*, which fall in the same lineage (Figure 2D). This suggests that heterochromatic mCHH is significantly reduced in many lineages of the Poaceae.

The methylome will be a composite of methylated and unmethylated regions. We implemented an approach (see Materials and methods) to identify methylated regions within a single sample to discern the average size of methylated regions and their level of DNA methylation for each species in each sequence context (**Figure S8**). For most species, regions of higher methylation are often smaller in size, with regions of low or intermediate methylation being larger (**Figure S9**). More small RNAs, in particular 24 nucleotide (nt) siRNAs map to regions of higher mCHH methylation (**Figure S10**). This may be because RdDM is primarily found on the edges of transposons whereas other mechanisms predominate in regions of deep heterochromatin (Zemach, Kim et al. 2013). Using these results, we can make inferences into the architecture of the methylome.

mCHG and mCHH regions are more variable in both size and methylation levels than mCG regions, as little variability in mCG regions was found between species (**Figure S8**). For mCHG regions, the Brassicaceae differed the most having lower methylation levels and *E. salsugineum* the lowest. This fits with *E. salsugineum* being a *cmt3* mutant and RdDM being responsible for residual mCHG (Bewick, Ji et al. 2016). However, the sizes of these regions are similar to other species, indicating that this has not resulted in fragmentation of these regions (**Figure S8**). The most variability was found in mCHH regions. Within the Fabaceae, the bulk of mCHH regions in *G. max* and *P. vulgaris* are of lower methylation in contrast to *M. truncatula* and *L. japonicus* (**Figure S8**). As these lower methylated mCHH regions are larger in size (Figure S9) and less targeted by 24 nt siRNAs (**Figure S10**), it would appear that deep heterochromatin mechanisms, like those mediated by CMT2, are more predominant than RdDM in these species as compared to *M. truncatula* and *L. japonicus.* Indeed the genomes of *G. max* and *P. vulgaris* are also larger than *M. truncatula* and *L. japonicus* (**Table S2**). In the Poaceae, we also find that mCHH regions are more highly methylated, even though genome-wide, mCHH levels are lower (**Figure S8**). This indicates that much of the mCHH in these genomes comes from smaller regions targeted by RdDM (**Figure S9, S10**), which is supported by RdDM mutants in *Z. mays* (Li, Eichten et al. 2014). In contrast, previously discussed species like *M. esculenta, T. cacao*, and *V. vinifera* had mCHH regions of both low methylation and small size which could indicate that effect of all mCHH pathways have been limited in these species (**Figure S9, S10**).

## Methylation of repeats

Genome-wide mCG and mCHG levels are related to the proliferation of repetitive elements. Although the quality of repeat annotations does vary between the species studied, correlations were found between repeat number and mCG (p-value=5 × 10^−5^) and mCHG levels (p-value=7.5 × 10^−3^) (Figure 3A, Table S3). This likely also explains the correlation of methylation with genome size, as large genomes often have more repetitive elements (Flavell, Bennett et al. 1974, Bennetzen, Ma et al. 2005). No such correlation between mCHH levels and repeat numbers was found after multiple testing correction (p-value > 0.05) (Figure 3A). This was unexpected given that mCHH is generally associated with repetitive sequences (Zemach, Li et al. 2005, Stroud, Do et al. 2014). Although coding sequence (CDS) mCHG and mCHH correlates with genome size (Figure 2B), only CDS mCHG correlated with the total number of repeats (p-value=3.8 × 10^−2^) (Figure S11A). All methylation contexts, however, were found to be correlated to the percentage of genes containing repeats within the gene (exons, introns, and untranslated regions - mCG p-value= 2.2 × 10^−3^, mCHG p-value=3.6 × 10^−4^, mCHH p-value = 2 × 10^−4^) (**Figure S11B, Table S3**), including introns or untranslated regions. In fact, plotting the percentage of genes containing repeats against the total number of repeats showed a cluster of species possessing fewer genic repeats given the total number of repeats in their genomes (**Figure S11C**). This implies that the transposon load of a genome alone does not affect methylation levels in genes, rather, it is more likely a result of the distribution of transposons within a genome.

**Fig. 3.**
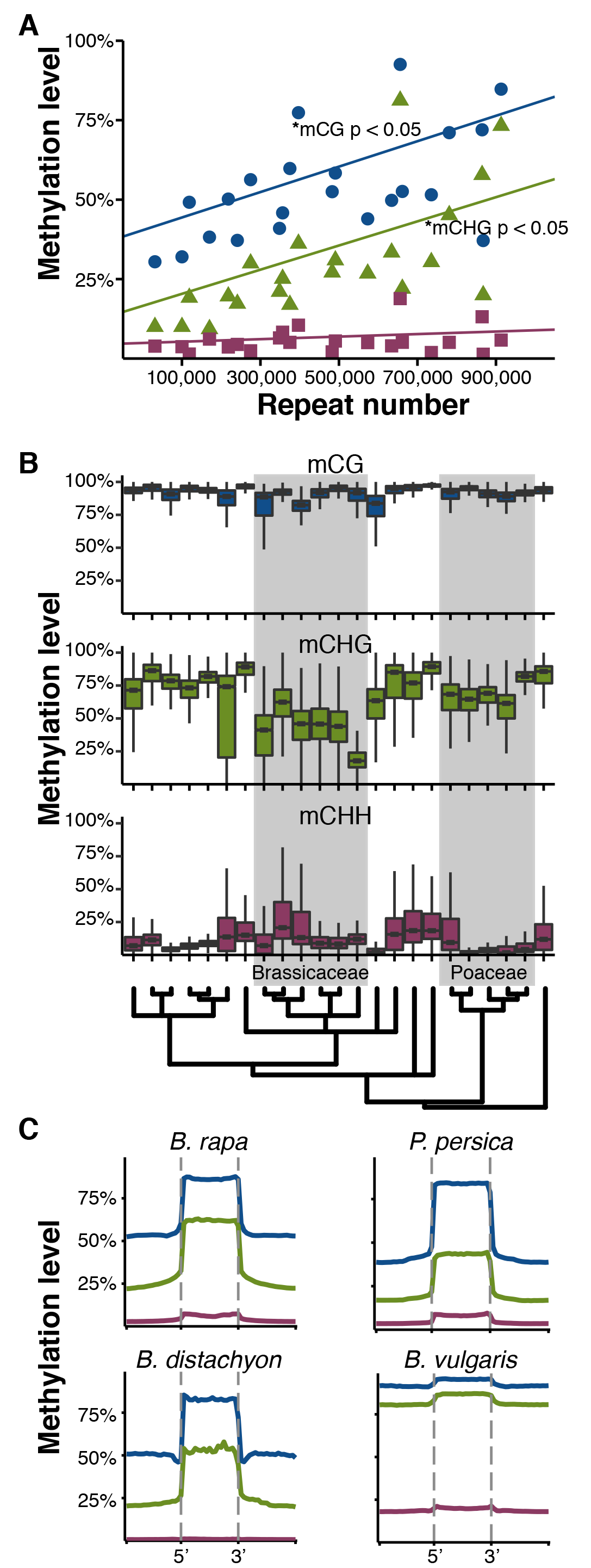
**(A)** Genome-wide methylation levels were correlated with repeat number for mCG (blue) and mCHG (green), but not for mCHH (maroon). Significant relationships indicated. **(B)** Distribution of methylation levels for repeats in each species. **(C)** Patterns of methylation upstream, across, and downstream of repeats for mCG (blue), mCHG (green), and mCHH (maroon).

Considerable variation exists in methylation patterns within repeats. Across all species, repeats were heavily methylated at CG sequences, but were more variable in CHG and CHH methylation (Figure 3B). mCHG was typically high at repeats in most species, with the exception of the Brassicaceae, in particular *E. salsugineum.* Similarly, low levels of mCHH were found in most Poaceae. Across the body of the repeat, most species show elevated levels in all three methylation contexts as compared to outside the repeat (Figure 3C, **S12**). Again, several Poaceae species stood out, as *B. distachyon* and *Z. mays* showed little change in mCHH within repeats, fitting with the observation that mCHH is depleted in deep heterochromatic regions of the Poaceae.

## CG gene body methylation

Methylation within genes in all three contexts is associated with suppressed gene expression (Law and Jacobsen 2010), whereas genes that are only mCG methylated within the gene body are often constitutively expressed genes (Tran, Henikoff et al. 2005, Zhang, Yazaki et al. 2006, Zilberman, Gehring et al. 2007). We classified genes using a modified version of the binomial test described by Takuno and Gaut (Takuno and Gaut 2012) into one of four categories: CG gene body methylated (hereafter gbM), mCHG, mCHH, and unmethylated (UM) (**Figure S13**, **Table S4**). gbM genes are methylated at CG sites, but not at CHG or CHH. NonCG contexts are often coincident with mCG, for example RdDM regions are methylated in all three contexts. We further classified nonCG methylated genes as mCHG genes (mCHG and mCG, no mCHH) or mCHH genes (mCHH, mCHG, and mCG). Genes with insignificant amounts of methylation were classified as unmethylated.

Between species, the methylation status of gbM can be conserved across orthologs (Takuno and Gaut 2013). The methylation state of orthologous genes across all species was compared using *A. thaliana* as an anchor (Figure 4A). *A. lyrata* and *C. rubella* are the most closely related to *A. thaliana* and also have the greatest conservation of methylation status, with many *A. thaliana* gbM gene orthologs also being gbM genes in these species (~86.3% and ~79.8% of *A. thaliana* gbM genes, respectively). However, they also had many gbM genes that had unmethylated *A. thaliana* orthologs (~ 18.6% and ~13.9% of *A. thaliana* genes, respectively). Although gbM is generally “conserved” between species, this conservation breaks down over evolutionary distance with gains and losses of gbM in different lineages. In terms of total number of gbM genes, *M. truncatula* and *Mimulus guttatus* had the greatest number (**Table S2**). However, when the percentage of gbM genes in the genome is taken into account (Figure 4B), *M. truncatula* appeared similar to other species, whereas *M. guttatus* remained an outlier with ~60.7% of all genes classified as gbM genes. The reason why *M. guttatus* has unusually large numbers of gbM loci is unknown and will require further investigation. In contrast, there has been considerable loss of gbM genes in *Brassica rapa*, and *Brassica oleracea* and a complete loss in *E. salsugineum.* This suggests that over longer evolutionary distance, the methylation status of gbM varies considerably and is dispensable as it is lost entirely in *E. salsugineum.*

**Fig. 4.**
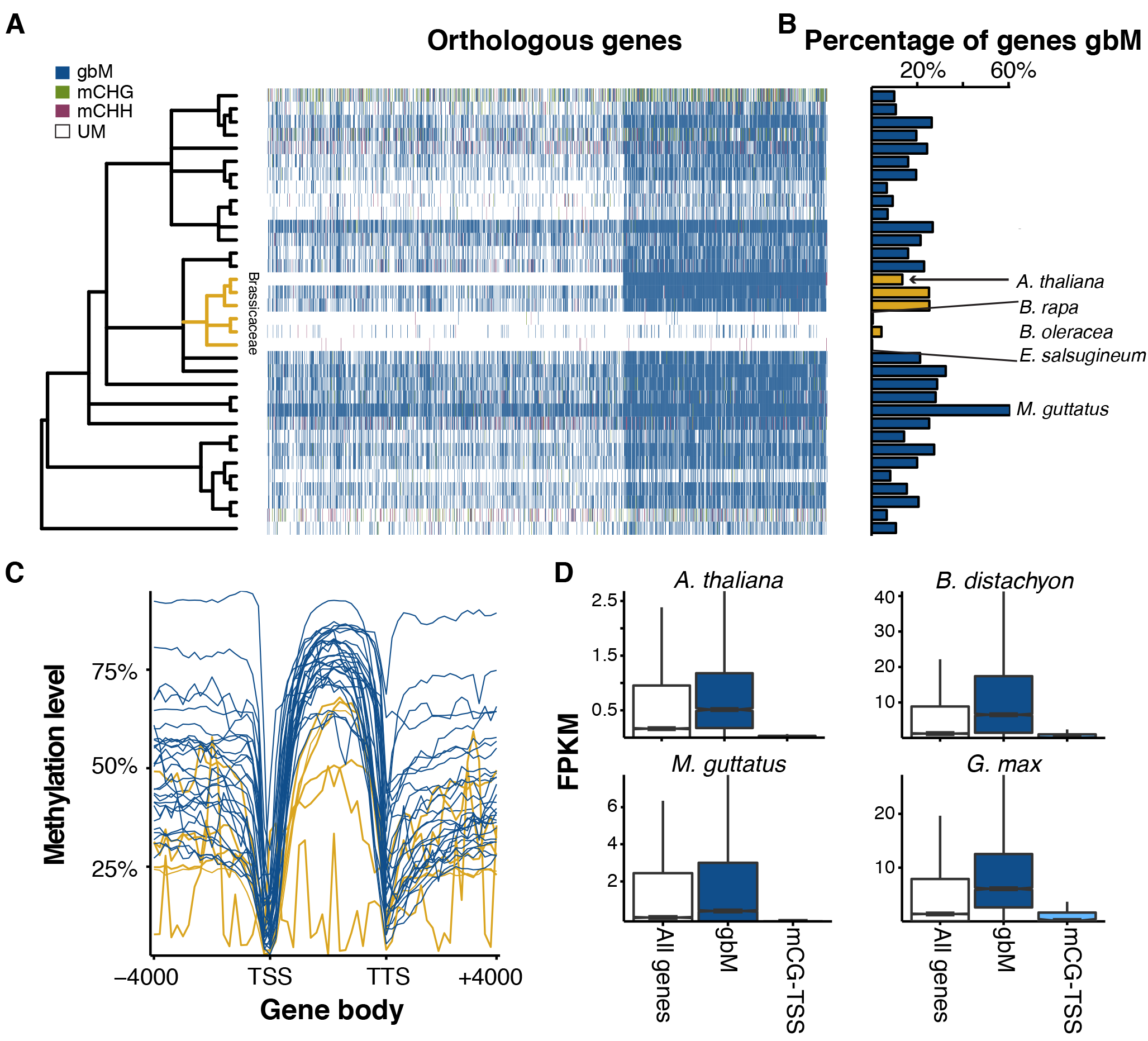
**(A)** Heatmap showing methylation state of orthologous genes (horizontal axis) to *A. thaliana* for each species (vertical axis). Species are organized according to phylogenetic relationship. **(B)** Percentage of genes in each species that are gbM (mCG only in coding sequences). The Brassicacea are highlighted in gold. **(C)** The levels of mCG in upstream, across, and downstream of gbM genes for all species. Species in gold belong to the Brassicaceae and illustrate the decreased levels and loss of mCG. **(D)** gbM genes are more highly expressed, while mCG over the TSS (mCG-TSS) has reduced gene expression.

GbM is characterized by a sharp decrease of methylation around the transcriptional start site (TSS), increasing mCG throughout the gene body, and a sharp decrease at the transcriptional termination site (TTS) (Zhang, Yazaki et al. 2006, Zilberman, Gehring et al. 2007). gbM genes identified in most species show this same trend and even have comparable levels of methylation (Figure 4C, **S14**). Here, too, the decay and loss of gbM in the Brassicaceae is observed as *B. rapa* and *B. oleracea* have the second and third lowest methylation levels, respectively in gbM genes and *E. salsugineum* shows no canonical gbM having only a few genes that passed statistical tests for having mCG in gene bodies. As has previously been found (Zhang, Yazaki et al. 2006, Zilberman, Gehring et al. 2007), gbM genes are more highly expressed as compared to UM and nonCG (mCHG and mCHH) genes (Figure 4D, **S15**). The exception to this is *E. salsugineum* where gbM genes have almost no expression. A subset of unexpressed genes with mCG methylation was found, and in some cases, had higher mCG methylation around the TSS (mCG-TSS). Using previously identified mCG regions we identified genes with mCG overlapping the TSS, but lacking either mCHG or mCHH regions within or near genes. These genes had suppressed expression (Figure 4D, **S15**) showing that although mCG is not repressive in gene-bodies, it can be when found around the TSS.

gbM genes are known to have many distinct features in comparison to UM genes. They are typically longer, have more exons, the observed number of CG dinucleotides in a gene are lower than expected given the GC content of the gene ([O/E]), and they evolve more slowly (Takuno and Gaut 2012, Takuno and Gaut 2013). We compared gbM genes to UM genes for each of these characteristics, using *A. thaliana* as the base for pairwise comparison for all species except the Poaceae where *O. sativa* was used (**Table S5**). With the exception of *E. salsugineum*, which lacks canonical gbM, these genes were longer and had more exons than UM genes (**Table S5**). Most gbM genes also had a lower CG [O/E] than UM genes, except for six species, four of which had a greater CG [O/E]. These included both *M. guttatus* and *M. truncatula*, which had the greatest number of gbM genes of any species. Recent conversion of previously UM genes to a gbM status could in part explain this effect. Previous studies have shown that gbM orthologs between *A. thaliana* and *A. lyrata* (Takuno and Gaut 2012) and between *B. distachyon* and *O. sativa* (Takuno and Gaut 2013) are more slowly evolving than UM orthologs. Although this result holds up over short evolutionary distances, it breaks down over greater distances with gbM genes typically evolving at equivalent rates as UM and, in some cases, faster rates (**Table S5**).

## NonCG methylated genes

NonCG methylation exists within genes and is known to suppress gene expression (Bender and Fink 1995, Cubas, Vincent et al. 1999, Manning, Tor et al. 2006, Martin, Troadec et al. 2009, Durand, Bouche et al. 2012). Although differences in annotation quality could lead to some transposons being misannotated as genes and thus as targets of nonCG methylation, within-species epialleles demonstrates that significant numbers of genes are indeed targets (Schmitz, He et al. 2013, Schmitz, Schultz et al. 2013). In many species there were genes with significant amounts of mCHG and little to no mCHH. High levels of mCHG within *Z. mays* genes is known to occur, especially in intronic sequences due in part to the presence of transposons (West, Li et al. 2014). Based on this difference in methylation, mCHG and mCHH genes were maintained as separate categories (**Table S4**). The methylation profiles of mCHG and mCHH genes often resembled that of repeats (Figure 5A, **S14**). Both mCHG and mCHH genes are associated with reduced expression levels (Figure 5B, **S15**). As mCHG methylation is present in mCHH genes, this may indicate that mCHG alone is sufficient for reduced gene expression. It was also observed that *Cucumis sativus* has an unusual pattern of mCHH in many highly expressed genes. This pattern was not observed in a second *C. sativus* sample and will require further study to understand its cause (**Figure S16**). The number of genes possessing nonCG types of methylation ranged from as low as ~3% of genes (*M. esculenta*) to as high as ~32% of genes (*F. vesca)* (Figure 5C). In all the Poaceae, mCHG genes made up at least ~5% of genes and typically more. In contrast, mCHG genes were relatively rare in the Brassicaceae where mCHH genes were the predominant type of nonCG genes. In most of the clonally propagated species with low mCHH, there were typically few mCHH genes and more mCHG genes with the exception of *F. vesca.*

**Fig. 5.**
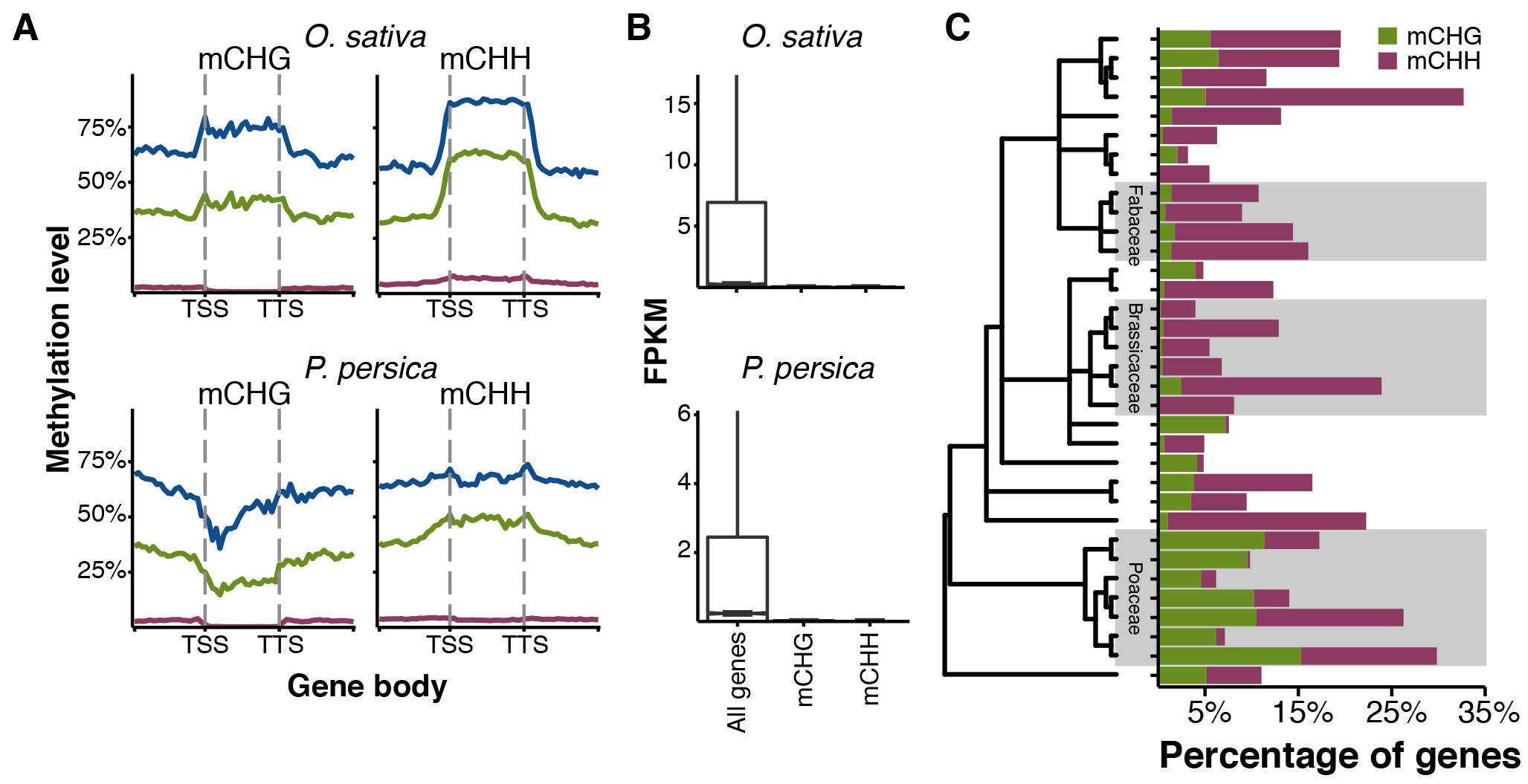
**(A)** Methylation levels for mCG (blue) (mCG only in coding sequences), mCHG (green) (mCG and mCHG in coding sequences), and mCHH (maroon) (mCG, mCHG, and mCHH in coding sequences) were plotted upstream, across, and downstream of mCHG and mCHH genes. **(B)** Gene expression of mCHG and mCHH genes versus all genes. **(C)** The percentage of mCHG and mCHH genes per species. Species are arranged by phylogenetic relationship.

Unlike gbM genes, there was no conservation of methylation status across orthologs of mCHG and mCHH genes (**Figure S17**). For many nonCG methylated genes, orthologs were not identified based on our approach of reciprocal best BLAST hit. For example, orthologs were found for only 488 of 999 of *A. thaliana* mCHH genes across all species. Previous comparisons of *A. thaliana, A. lyrata*, and *C. rubella* have shown no conservation of nonCG methylation between orthologs within the Brassicaceae (Seymour, Koenig et al. 2014). However, we did observe some conservation based on gene ontology (GO). The same GO terms were often enriched in multiple species (**Figure S18**, **Table S6**). The most commonly enriched terms were involved in processes such as proteolysis, cell death, and defense responses; processes that could have profound effects on normal growth and development and may be developmentally or environmentally regulated. There was also enrichment in many species for genes related to electron-transport chain processes, photosynthetic activity, and other metabolic processes. Further investigation of these genes revealed that many are orthologs to chloroplast or mitochondrial genes, suggesting that they may be recent transfers from the organellar genome. The transfer of organellar genes to the nucleus is a frequent and ongoing process (Stegemann, Hartmann et al. 2003, Roark, Hui et al. 2010). Although DNA methylation is not found in chloroplast genomes, transfer to the nucleus places them in a context where they can be methylated, contributing to the mutational decay of these genes via deamination of methylated cytosines (Huang, Grunheit et al. 2005).

## mCHH islands

In Z. *mays*, high mCHH is enriched in the upstream and downstream regions of highly expressed genes and are termed mCHH islands (Gent, Ellis et al. 2013, Li, Gent et al. 2015). We identified mCHH islands 2kb upstream and downstream of annotated genes for each species, finding that the percentage of genes with such regions varied considerably across species (Figure 6A) the fewest being in *V. vinifera, B. oleracea*, and *T. cacao*, each having mCHH islands associated with less than 2% of genes. As both *V. vinifera* and *T.* cacao have low genome-wide mCHH levels, this may explain the difference, however, this is not the case for *B. oleracea.* mCHH islands are thought to mark euchromatin-heterochromatin boundaries and are often associated with transposons, however, we found no correlation between the total number of repeats in the genome and the number of genes with mCHH islands (**Figure S19A**, **Table S3**). As in the case of CDS methylation levels, this lack of correlation was largely due to differences in the distribution of repeats (**Figure S19B**, **Table S3**). When correlated to the percentage of genes with repeats 2kb upstream or downstream, both upstream and downstream mCHH islands are correlated (upstream p-value = 7.9 × 10^−6^, downstream p-value < 8.1 × 10^−6^) (Figure 6B). *B. oleracea* in particular stood out as having few repeats in 2kb upstream or downstream of genes, explaining in part why it possess so few mCHH islands, despite its large genome and overall number of repeats.

**Fig. 6.**
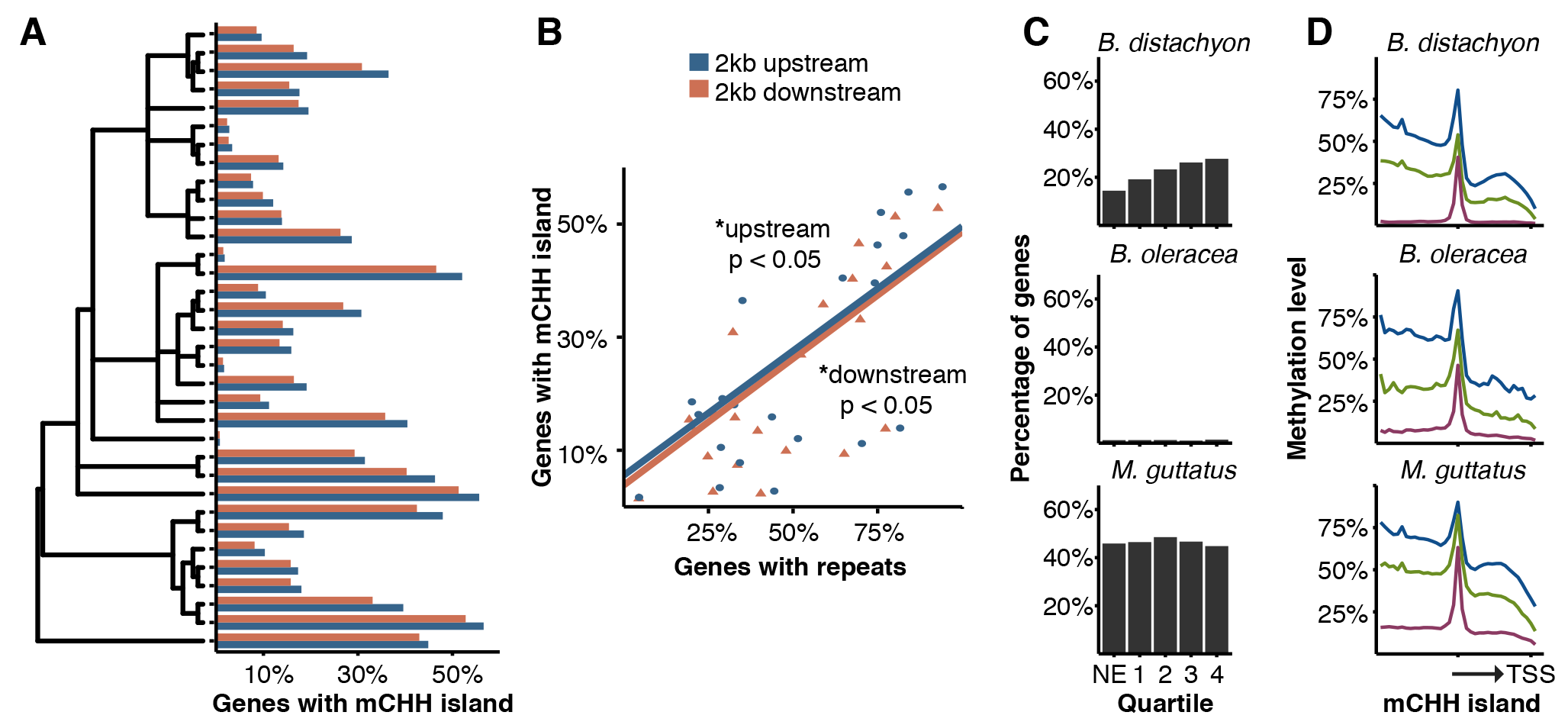
**(A)** Percentage of genes with mCHH islands 2kb upstream or downstream. **(B)** Upstream and downstream mCHH islands are correlated with upstream and downstream repeats (respectively). Significant relationships indicated. **(C)** Association of upstream mCHH islands with gene expression. Genes are divided into non-expressed (NE) and quartiles of increasing expression. **(D)** Patterns of upstream mCHH islands. Blue, green, and red lines represent mCG, mCHG and mCHH levels, respectively.

In *Z. mays*, mCHH islands are more common in genes in the most highly expressed quartiles (Gent, Ellis et al. 2013, Li, Gent et al. 2015). This is true of several species, such as *P. persica* and all the Poaceae (Figure 6C, **S20**). However, many species showed no significant association between mCHH islands and gene expression (Figure 6C, **S20**). This was true for all the Brassicaceae. Other species such as *M. guttatus*, despite having high levels of mCHH islands associated with genes like the Poaceae (~46% upstream, ~40% downstream), also showed no association with gene expression (Figure 6C). As has been observed previously in *Z. mays*, mCG and mCHG levels are generally higher on the distal side of the mCHH island to the gene (Figure 6D, **S21**), marking a boundary of euchromatin and heterochromatin (Li, Gent et al. 2015). However, this difference in methylation level is much less pronounced in most other species as compared to *Z. mays* and much less evident for downstream mCHH islands than for upstream ones (**Figure S21**). These may indicate different preferences in transposon insertion sites or a need to maintain a boundary of heterochromatin near the start of transcription.

## Discussion

We present the methylomes of 34 different angiosperm species in a phylogenetic framework using comparative epigenomics, which enables the study of DNA methylation in an evolutionary context. Extensive variation was found between species, both in levels of methylation and distribution of methylation, with the greatest variation being observed in nonCG contexts. The Brassicaceae show overall reduced mCHG levels and reduced numbers of gbM genes, leading to a complete loss in *E. salsugineum*, that is associated with loss of CMT3 (Bewick, Ji et al. 2016). Whereas in the Poaceae, mCHH levels are typically lower than that in other species. The Poaceae have a distinct epigenomic architecture compared to eudicots, with mCHH often depleted in deep heterochromatin and enriched in genic regions. We also observed that many species with a history of clonal propagation have lower mCHH levels. Epigenetic variation induced by propagation techniques can be of agricultural and economic importance (Ong-Abdullah, Ordway et al. 2015), and understanding the effects of clonal propagation will require future studies over multiple generations. Evaluation of per-site methylation levels, methylated regions, their structure, and association with small RNAs indicates that there are differences in the predominance of various molecular pathways.

Variation exists within features of the genome. Repeats and transposons show variation in their methylation level and distribution with impacts on methylation within genes and regulatory regions. Although gbM genes do show many conserved features, this breaks down with increasing evolutionary distance and as gbM is gained or lost in some species. gbM is known to be absent in the basal plant species *Marchantia polymorpha* (Takuno, Ran et al. 2016) and *Selaginella moellendorffii* (Zemach, McDaniel et al. 2010). That it has also been lost in the angiosperm *E. salsugineum*, indicates that it is dispensable over evolutionary time. That nonCG methylation shows no conservation at the level of individual genes, indicates that it is gained and lost in a lineage specific manner. It is an open question as to the evolutionary origins of nonCG methylation within genes. That these are correlated to repetitive elements in genes suggests transposons as one possible factor. That many nonCG genes lack orthologous genes could indicate a preferential targeting of *de novo* genes, as in the case of the QQS gene in *A. thaliana* (Silveira, Trontin et al. 2013). At a higher order level, there appears to be a commonality in what categories of genes are targeted, as many of the similar functions are enriched across species. Other features, such as mCHH islands, also are not conserved and show extensive variation that is associated with the distribution of repeats upstream and downstream of genes.

This study demonstrates that widespread variation in methylation exists between flowering plant species. For many species, this is the first reported methylome and methylome browsers for each species have been made available to serve as a resource (http://schmitzlab.genetics.uga.edu/plantmethylomes). Historically, our understanding has come primarily from *A. thaliana*, which has served as a great model for studying the mechanistic nature of DNA methylation. However, the extent of variation observed previously (Seymour, Koenig et al. 2014, Takuno, Ran et al. 2016) and now shows that there is still much to be learned about underlying causes of variation in this molecular trait. Due to its role in gene expression and its potential to vary independently of genetic variation, understanding these causes will be necessary to a more complete understanding of the role of DNA methylation underlying biological diversity.

## Materials and methods

### MethylC-seq and analysis

In plants, DNA methylation is highly stable between tissues and across generations (Schmitz, Schultz et al. 2011, Seymour, Koenig et al. 2014), showing little variation between replicates. DNA was isolated from leaf tissue and MethylC-seq libraries for each species were prepared as previously described (Urich, Nery et al. 2015). Previously published datasets were obtained from public databases and reanalyzed (Tuskan, Difazio et al. 2006, Schmitz, Schultz et al. 2011, Amborella Genome 2013, Gent, Ellis et al. 2013, Schmitz, He et al. 2013, Stroud, Ding et al. 2013, Zhong, Fei et al. 2013, Seymour, Koenig et al. 2014, Bewick, Ji et al. 2016). Genome sequences and annotations for most species were downloaded from Phytozome 10.1 (http://phytozome.jgi.doe.gov/pz/portal.html) (Goodstein, Shu et al. 2012). The *L. japonicus* genome was downloaded from the *Lotus japonicus* Sequencing Project (http://www.kazusa.or.jp/lotus/) (Sato, Nakamura et al. 2008), the *B. vulgaris* genome was downloaded from *Beta vulgaris* Resource bvseq.molaen.mpg.de/) (Dohm, Minoche et al. 2014), and the *C. sativa* genome from *C. sativa (Cannabis)* Genome Browser Gateway (http://genome.ccbr.utoronto.ca/cgi-bin/hgGatewav) (van Bakel, Stout et al. 2011). As annotations for *S. viridis* were not available, gene models from the closely related *S. italica* were mapped onto the *S. viridis* genome using Exonerate (Slater and Birney 2005) and the best hits retained.

Sequencing data for each species was aligned to their respective genome (Supplementary Table 1) (Arabidopsis Genome 2000, Jaillon, Aury et al. 2007, Ouyang, Zhu et al. 2007, Sato, Nakamura et al. 2008, Paterson, Bowers et al. 2009, Schnable, Ware et al. 2009, Chan, Crabtree et al. 2010, International Brachypodium 2010, Schmutz, Cannon et al. 2010, Velasco, Zharkikh et al. 2010, Hu, Pattyn et al. 2011, Shulaev, Sargent et al. 2011, van Bakel, Stout et al. 2011, Young, Debelle et al. 2011, Goodstein, Shu et al. 2012, Paterson, Wendel et al. 2012, Prochnik, Marri et al. 2012, Tomato Genome 2012, Amborella Genome 2013, Hellsten, Wright et al. 2013, International Peach Genome, Verde et al. 2013, Motamayor, Mockaitis et al. 2013, Slotte, Hazzouri et al. 2013, Yang, Jarvis et al. 2013, Dohm, Minoche et al. 2014, Myburg, Grattapaglia et al. 2014, Parkin, Koh et al. 2014, Schmutz, McClean et al. 2014, Wu, Prochnik et al. 2014) and methylated sites called using previously described methods (Schultz, He et al. 2015). In brief, reads were trimmed for adapters and quality using Cutadapt (Martin 2011) and then mapped to both a converted forward strand (all cytosines to thymines) and converted reverse strand (all guanines to adenines) using bowtie (Langmead 2010). Reads that mapped to multiple locations and clonal reads were removed. The non-conversion rate (rate at which unmethylated cytosines failed to be converted to uracil) was calculated by using reads mapping to the lambda genome or the chloroplast genome if available (Supplementary Table 1). Cytosines were called as methylated using a binomial test using the non-conversion rate as the expected probability followed by multiple testing correction using Benjamini-Hochberg False Discovery Rate (FDR). A minimum of three reads mapping to a site was required to call a site as methylated. Data are available at the Plant Methylome DB http://schmitzlab.genetics.uga.edu/plantmethylomes.

### Phylogenetic Tree

A species tree was constructed using BEAST2 (Bouckaert, Heled et al. 2014) on a set of 50 previously identified single copy loci (Duarte, Wall et al. 2010). Protein sequences were aligned using PASTA (Mirarab, Nguyen et al. 2015) and converted into codon alignments using custom Perl scripts. Gblocks (Castresana 2000) was used to identify conserved stretches of amino acids and then passed to JModelTest2 (Guindon and Gascuel 2003, Darriba, Taboada et al. 2012) to assign the most likely nucleotide substitution model.

### Genome-wide analyses

Genome-wide weighted methylation was calculated from all aligned data by dividing the total number of aligned methylated reads to the genome by the total number of methylated plus unmethylated reads (Schultz, Schmitz et al. 2012). To determine per-site methylation levels, the weighted methylation for each cystosine with at least 3 reads of coverage was calculated and this distribution plotted. Symmetry plots were constructed by identifying paired symmetrical cytosines that had sequencing coverage and plotting the per-site methylation level of the cytosine on the Watson strand against the per-site methylation level of the Crick strand. An *A. thaliana cmt3* mutant was used to empirically determine the per-site methylation level at which symmetrical methylation disappeared (Bewick, Ji et al. 2016) as 40%. Methylated symmetrical pairs above this level were considered to be symmetrically methylated, while those below as asymmetrical. Correlations between methylation levels, genome sizes, and gene numbers were done in R and corrected for phylogenetic signal using the APE (Paradis,Claude et al. 2004), phytools (Revell 2012), and NLME packages assuming a model of Brownian motion. In total, 22 comparisons were conducted (Supplementary Table 3) and a p-value < 0.05 after Bonferroni Correction. Distribution of methylation levels and genes across chromosomes was conducted by dividing the genome into 100 kb windows, sliding every 50 kb using BedTools (Quinlan and Hall 2010) and custom scripts. Pearson’s correlation between gene number and methylation level in each window was conducted in R. Weighted methylation levels for each repeat were calculated using custom python and R scripts.

### Methylated-regions

Methylated regions were defined independent of genomic feature by methylation context (CG, CHG, or CHH) using BEDTools (Quinlan and Hall 2010) and custom scripts. For each context, only methylated sites in that respective context were considered used to define the region. The genome was divided into 25bp windows and all windows that contained at least one methylated cytosine in the context of interest were retained. 25bp windows were then merged if they were within 100 bp of each other, otherwise they were kept separate. The merged windows were then refined so that the first methylated cytosine became the new start position and the last methylated cytosine new end position. Number of methylated sites and methylation levels for that region was then recalculated for the refined regions. A region was retained if it contained at least five methylated cytosines and then split into one of four groups based on the methylation levels of that region: group 1, < 0.05%, group 2, 5-15%, group 3, 15-25%, group 4, > 25%. Size of methylated regions were determined using BedTools.

### Small RNA (sRNA) cleaning and filtering

Libraries for *B. distachyon, C. sativus, E. grandis, E. salsugineum, M. truncatula, P. hallii*, and *R. communis* were constructed using the TruSeq Small RNA Library Preparation Kit (Illumina Inc). Small RNA-seq datasets for additional species were downloaded from GEO and the SRA and reanalyzed (Schmitz, Schultz et al. 2011, Amborella Genome 2013, International Peach Genome, Verde et al. 2013, Stroud, Ding et al. 2013, Chavez Montes, de Fatima Rosas-Cardenas et al. 2014, Wang, Yuan et al. 2015). The small RNA toolkit from the UEA computational Biology lab was used to trim and clean the reads (Stocks, Moxon et al. 2012). For trimming, 8 bp of the 3’ adapter was trimmed. Trimmed and cleaned reads were aligned using PatMan allowing for zero mismatches (Prufer, Stenzel et al. 2008). BedTools (Quinlan and Hall 2010) and custom scripts were used to calculate overlap with mCHH regions.

### Gene-level analyses

Genes were classified as mCG, mCHG, or mCHH by applying a binomial test to the number of methylated sites in a gene (Takuno and Gaut 2012) (**Figure S13**, **Table S4**). The total number of cytosines and the methylated cytosines were counted for each context for the coding sequences (CDS) of the primary transcript for each gene. A single expected methylation rate was estimated for all species by calculating the percentage of methylated sites for each context from all sites in all coding regions from all species. We restricted the expected methylation rate to only coding sequences as the species study differ greatly in genome size, repeat content, and other factors that impact genome-wide methylation. Furthermore, it is known that some species have an abundance of transposons in UTRs and intronic sequences, which could lead to misclassification of a gene. A single value was calculated for all species to facilitate comparisons between species and to prevent setting the expected methylation level to low, as in the case of *E. salsugineum* or to high, as in the case of *B. vulgaris*, which would further lead to misclassifications.

A binomial test was then applied to each gene for each sequence context and q-values calculated by adjusting p-values by Benjamini-Hochberg FDR. Genes were classified as mCG if they had reads mapping to at least 20 CG sites and has q-value < 0.05 for mCG and a q-value > 0.05 for mCHG and mCHH. Genes were classified as mCHG if they had reads mapping to at least 20 CHGs, a mCHG q-value < 0.05, and a mCHH q-value > 0.05. As mCG is commonly associated with mCHG, the q-value for mCG was allowed to be significant or insignificant in mCHG genes. Genes were classified as mCHH if they had reads mapping to at least 20 mCHH sites and a mCHH q-value < 0.05. Q-values for mCG and mCHG were allowed to be anything as both types of methylation are associated with mCHH. mCG-TSS genes were identified by overlap of mCG regions with the TSS of each gene and the absence of any mCHG or mCHH regions within the gene or 1000 bp upstream or downstream.

GO terms for each gene were downloaded from phytozome 10. 1 (http://phytozome.jgi.doe.gov/pz/portal.html) (Goodstein, Shu et al. 2012). GO term enrichment was performed using the parentCHILD algorithm (Grossmann, Bauer et al. 2007) with the F-statistic as implemented in the topGO module in R. GO terms were considered significant with a q-value < 0.05.

## Exon number, gene length and [O/E]

For each species the general feature format 3 (gff3) file from phytozome 10.1 (Goodstein, Shu et al. 2012) was used to determine exon number and coding sequence length (base pairs, bp) for each annotated gene (hereafter referred to as CDS). Additionally, for each full length CDS (starting with the start codon ATG, and ending with one of the three stop codons TAA/TGA/TAG), from the phytozome 10.1 (Goodstein, Shu et al. 2012) primary CDS fasta file, the CG [O/E] ratio was calculated, which is the observed number of CG dinucleotides relative to that expected given the overall G+C content of a gene. Differences for these genic features between CG gbM and UM genes were assessed using permutation tests (100,000 replicates) in R, with the null hypothesis being no difference between the gbM and UM methylated genes.

## Identifying orthologs and estimating evolutionary rates

Substitution rates were calculated between CDS pairs of monocots to *Oryza sativa*, and dicots to *A. thaliana.* Reciprocal best BLAST with an e-value cutoff of ≤1E-08 was used to identify orthologs between dicot-A. *thaliana*, and monocot-O. *sativa* pairs. Individual CDS pairs were aligned using MUSCLE (Edgar 2004), insertion-deletion (indel) sites were removed from both sequences, and the remaining sequence fragments were shifted into frame and concatenated into a contiguous sequence. A ≥30 bp and ≥300 bp cutoff for retained fragment length after indel removal, and concatenated sequence length was implemented, respectively. Coding sequence pairs were separated into each combination of methylation (i.e., CG gbM-CG gbM, and UM-UM). The *yn00* (Yang-Neilson) (Yang and Nielsen 2000) model in the program PAML for pairwise sequence comparison was used to estimate synonymous and non-synonymous substitution rates, and adaptive evolution (dS, *dN*, and ω, respectively) (Yang 1997). Differences in rates of evolution between methylated and unmethylated pairs were assessed using permutation tests (100,000 replicates) in R, with the null hypothesis being no difference between the CG gbM and UM methylated genes.

## RNA-seq mapping and analysis

RNA-seq datasets (Schmitz, Schultz et al. 2011, Brown, Kroon et al. 2012, Chodavarapu, Feng et al. 2012, Perazzolli, Moretto et al. 2012, Tomato Genome 2012, Amborella Genome 2013, International Peach Genome, Verde et al. 2013, Schmitz, He et al. 2013, Seymour, Koenig et al. 2014, Tang, Dong et al. 2014, Livingstone, Royaert et al. 2015, Wang, Yuan et al. 2015, Bewick, Ji et al. 2016) were downloaded from the Gene Expression Omnibus (GEO) and the NCBI Short Read Archive (SRA) for reanalysis. *B. distachyon* and *C. sativus* RNA-seq libraries were constructed using Illumina TruSeq Stranded mRNA Library Preparation Kit (Illumina Inc). and sequenced on a NextSeq500 at the Georgia Genomics Facility. Reads were aligned using Tophat v2.0.13 (Kim, Pertea et al. 2013) supplied with a reference genome feature file (GFF) with the following arguments-I 50000-b2-very-sensitive-b2-D 50. Transcripts were then quantified using Cufflinks v2.2.1 (Trapnell, Roberts et al. 2012) supplied with a reference GFF.

### mCHH islands

mCHH islands were identified for both upstream and downstream regions as previously described (Li, Gent et al. 2015). Briefly, methylation levels were determined for 100bp windows across the genome. Windows of 25% or greater mCHH either 2 kb upstream or downstream of genes were identified and intersecting windows of that region were retained. Methylation levels were then plotted centered on the window of highest mCHH. Genes associated with mCHH islands were categorized as nonexpressed (NE) or divided into one of four quartiles based on their expression level.

## Acknowledgements

We would like to thank Drs. J. Chris Pires, Scott T. Woody, Richard M. Amasino, Heinz Himmelbauer, Fred G. Gmitter, Timothy R. Hughes, Rebecca Grumet, CJ Tsai, Karen S. Schumaker, Kevin M. Folta, Marc Libault, Steve van Nocker, Steve D. Rounsely, Andrea L. Sweigart, Gerald A. Tuskan, Thomas E. Juenger, Douglas G. Bielenberg, Brian Dilkes, Thomas P. Brutnell, Todd C. Mockler, Mark J. Guiltinan, and Mallikarjuna K. Aradhya for providing tissue and DNA of various species used in this study. The work conducted by the U.S. Department of Energy Joint Genome Institute is supported by the Office of Science of the U.S. Department of Energy under Contract No. DE-AC02-05CH11231 to J.S. We thank the Joint Genome Institute and collaborators for access to unpublished genomes of *B. rapa, S. viridis, P. virgatum*, and *P. hallii*. This work was supported by the National Science Foundation (NSF) (MCB-1402183), by the Office of the Vice President of Research at UGA, and by The Pew Charitable Trusts to R.J.S. C.E.N was supported by a NSF postdoctoral fellowship (IOS-1402183).

## Author information

Genome browsers for all methylation data used in this paper is located at Plant Methylation DB (http://schmitzlab.genetics.uga.edu/plantmethylomes). Sequence data for MethylC-seq, RNA-seq, and small RNA-seq is located at the Gene Expression Omnibus, accession GSE79526.

## Author Contributions

Conceptualization, C.E.N., A.J.B.and R.J.S.; Performed Experiments, C.E.N., N.A.R., K.D.K., A.R., J.T.P., and R.J.S.; Data Analysis, C.E.N., A.J.B., L.J., M.S.A.; Writing - Original Draft, C.E.N; Writing - Review & Editing, C.E.N., A.J.B,. S.A.J., N.M.S., and R.J.S.; Resources, Q.L., J.M.B., J.A.U., C.E., J.S., J.G., S.A.J., N.M.S.

